## Supplemental Figures for "Disruption of the CRF_1_ receptor eliminates morphine-induced sociability deficits and firing of oxytocinergic neurons in male mice"

Manuscript title:

List of the supplementary figures:

**Fig. S1. Locomotor activity of C57BL/6J mice during the three-chamber test with morphine**.

**Fig. S2. Locomotor activity of $CRF_1$ receptor-deficient mice during the three-chamber test with morphine**.

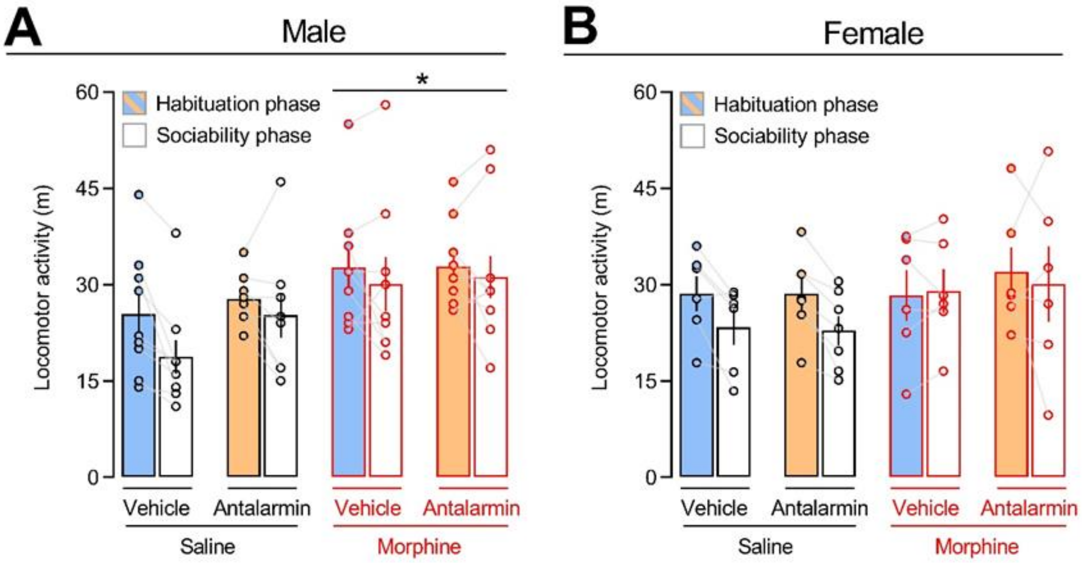


**Fig. S1. Locomotor activity of C57BL/6J mice during the three-chamber test with morphine**. Distance (m) travelled by (**A**) male and (**B**) female C57BL/6J mice treated with either vehicle or antalarmin (20 mg/kg, p.o.) followed by either saline or morphine (2.5 mg/kg, i.p.) during the habituation and the sociability phases of the three-chamber test. Overall, male ($P<0.005$) and female ($P<0.05$) mice travelled more distance during the habituation than during the sociability phase. N=8-10/group for male mice; n=6-7/group for female mice. The number of animals within each experimental group is reported in **Table S1A**. Values represent mean±SEM. *$P<0.05$ *vs.* saline-treated mice, independently of vehicle or antalarmin treatment.

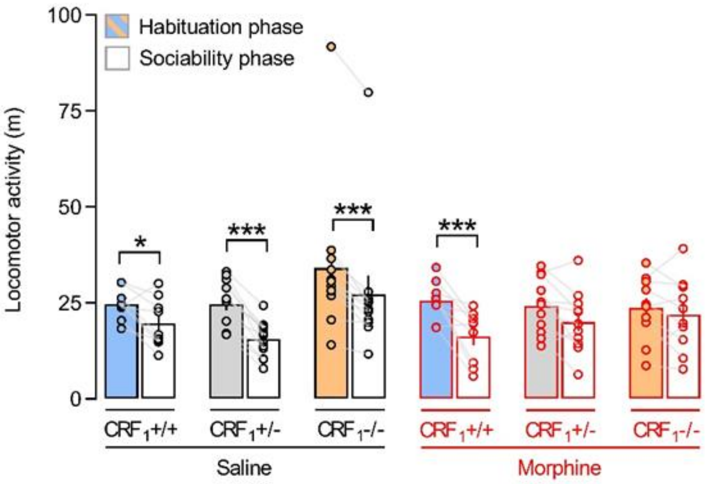


**Fig. S2. Locomotor activity of $CRF_1$ receptor-deficient mice during the three-chamber test with morphine**. Distance (m) travelled by saline- or morphine (0.625 mg/kg, i.p.)-treated male $CRF_1$+/+, $CRF_1$+/- and $CRF_1$-/- mice during the habituation and the sociability phases of the three-chamber test. N=9-13/group. The number of animals within each experimental group is reported in **Table S1B**. Values represent mean±SEM. *P<0.05, ***P<0.0005.
