## Supplemental Tables for "Disruption of the CRF_1_ receptor eliminates morphine-induced sociability deficits and firing of oxytocinergic neurons in male mice"

Manuscript title:

List of the supplementary tables:

**Table S1. Number of animals used and cells patched and recorded**.

**Table S2. Statistical analysis of the three-chamber sociability test in C57BL/6J mice**.

**Table S3. Statistical analysis of locomotor activity displayed by C57BL/6J mice during the three-chamber sociability test**.

**Table S4. Statistical analysis of the three-chamber sociability test in $CRF_1$ receptor-deficient mice**.

**Table S5. Female $CRF_1$+/+ and $CRF_1$+/- mice fail to perform in the three-chamber task for sociability**.

**Table S6. Statistical analysis of neuronal firing in C57BL/6J mice**.

**Table S1. Number of animals used and cells patched and recorded**. **A**) Number of male and female C57BL/6J mice tested in the three-chamber task after being treated *per os* (p.o.) with either vehicle (veh) or antalarmin (anta; 20 mg/kg) and intraperitoneally (i.p.) with either saline (sal) or morphine (mor; 2.5 mg/kg). **B**) Number of saline- or morphine (0.625 mg/kg)-treated male $CRF_1$+/+, $CRF_1$+/- and $CRF_1$-/- mice tested in the three-chamber task and number of cells patched and recorded in the electrophysiology studies. In brackets, the number of animals excluded from the statistical analysis of the data because met the exclusion criterion, i.e., exploration of each region of interest (ROI, side half-chamber) of the three-chamber apparatus for more than 80%, or less than 20%, of the total time spent in both ROIs during the habituation phase. **C**) Number of patched and recorded paraventricular nucleus of the hypothalamus (PVN) neurons expressing oxytocin (OXY) and/or arginine-vasopressin (AVP) or neither OXY nor AVP (no staining) in male (M) and female (F) C57BL/6J mice treated with either vehicle or antalarmin followed by either saline or morphine, as in **A**.

**A**

| **C57BL/6J** | **Veh/sal** | **Anta/sal** | **Veh/mor** | **Anta/mor** |
|---|---|---|---|---|
| **Male** | 9 (0) | 9 (1) | 12 (3) | 12 (2) |
| **Female** | 7 (1) | 8 (1) | 7 (1) | 9 (3) |

**B**

| | **Three-chamber** | | **Electrophysiology** | |
|---|---|---|---|---|
| **Genotype** | **Sal** | **Mor** | **Sal** | **Mor** |
| **$CRF_1$+/+** | 12 (3) | 11 (2) | 18 | 21 |
| **$CRF_1$+/-** | 16 (3) | 17 (6) | 21 | 31 |
| **$CRF_1$-/-** | 13 (1) | 12 (2) | 17 | 17 |

**C**

| | **Veh/sal** | | **Anta/sal** | | **Veh/mor** | | **Anta/mor** | |
|---|---|---|---|---|---|---|---|---|
| **C57BL/6J** | **M** | **F** | **M** | **F** | **M** | **F** | **M** | **F** |
| **OXY/AVP** | 7 | 7 | 8 | 7 | 11 | 8 | 14 | 9 |
| **AVP** | 13 | 2 | 16 | 2 | 12 | 2 | 8 | 1 |
| **OXY** | 2 | 11 | 1 | 7 | 1 | 8 | 2 | 12 |
| **No staining** | 2 | 6 | 4 | 4 | 4 | 4 | 5 | 3 |
| ***Total* cells** | **24** | **26** | **29** | **20** | **28** | **22** | **29** | **25** |

**Table S2. Statistical analysis of the three-chamber sociability test in C57BL/6J mice**. Statistical analysis of time (s) spent in the regions of interest (ROIs, side half-chambers) of the three-chamber apparatus by male and female C57BL/6J mice treated *per os* (p.o.) with either vehicle or antalarmin (20 mg/kg) and intraperitoneally (i.p.) with either saline or morphine (2.5 mg/kg), during the habituation or the sociability phase of the three-chamber test. Pretreatment (P): vehicle *vs*. antalarmin. Treatment (T): saline *vs*. morphine. Repeated measures (RM): mouse *vs*. object. Further details are reported in the "statistical analysis" section of the manuscript.

| | **Male** | | **Female** | |
|---|---|---|---|---|
| | **Habituation** | **Sociability** | **Habituation** | **Sociability** |
| **P** | $F_{1,32}$=1.122 P=0.297 | $F_{1,32}$=4.075 P=0.052 | $F_{1,21}$=1.692 P=0.207 | $F_{1,21}$=1.296 P=0.268 |
| **T** | $F_{1,32}$=0.064 P=0.802 | $F_{1,32}$=0.438 P=0.513 | $F_{1,21}$=23.501 P<0.0001 | $F_{1,21}$=25.077 P<0.0001 |
| **P x T** | $F_{1,32}$=1.025 P=0.319 | $F_{1,32}$=11.590 P<0.005 | $F_{1,21}$=0.253 P=0.620 | $F_{1,21}$=0.290 P=0.596 |
| **RM** | $F_{1,32}$=0.001 P=0.976 | $F_{1,32}$=23.795 P<0.0001 | $F_{1,21}$=0.005 P=0.942 | $F_{1,21}$=4.547 P<0.05 |
| **P x RM** | $F_{1,32}$=0.128 P=0.723 | $F_{1,32}$=8.977 P<0.01 | $F_{1,21}$=0.024 P=0.879 | $F_{1,21}$=0.032 P=0.860 |
| **T x RM** | $F_{1,32}$=0.240 P=0.628 | $F_{1,32}$=1.225 P=0.277 | $F_{1,21}$=0.010 P=0.922 | $F_{1,21}$=14.218 P<0.005 |
| **P x T x RM** | $F_{1,32}$=0.163 P=0.689 | $F_{1,32}$=8.444 P<0.01 | $F_{1,21}$=0.002 P=0.965 | $F_{1,21}$=0.037 P=0.848 |

**Table S3. Statistical analysis of locomotor activity displayed by C57BL/6J mice during the three-chamber sociability test**. Statistical analysis of distance (m) travelled by male and female C57BL/6J mice treated *per os* (p.o.) with either vehicle or antalarmin (20 mg/kg) and intraperitoneally (i.p.) with either saline or morphine (2.5 mg/kg), during the habituation and the sociability phases of the three-chamber test. Pretreatment (P): vehicle *vs*. antalarmin. Treatment (T): saline *vs*. morphine. Repeated measures (RM): habituation *vs*. sociability phase. Further details are reported in the "statistical analysis" section of the manuscript.

| | **Male** | **Female** |
|---|---|---|
| **P** | $F_{1,32}$=0.799 P=0.378 | $F_{1,21}$=0.107 P=0.746 |
| **T** | $F_{1,32}$=6.648 P<0.05 | $F_{1,21}$=1.485 P=0.237 |
| **P x T** | $F_{1,32}$=0.401 P=0.531 | $F_{1,21}$=0.159 P=0.694 |
| **RM** | $F_{1,32}$=9.779 P<0.005 | $F_{1,21}$=6.055 P<0.05 |
| **P x RM** | $F_{1,32}$=1.708 P=0.201 | $F_{1,21}$=0.431 P=0.519 |
| **T x RM** | $F_{1,32}$=1.431 P=0.240 | $F_{1,21}$=3.929 P=0.061 |
| **P x T x RM** | $F_{1,32}$=0.482 P=0.492 | $F_{1,21}$=0.143 P=0.709 |

**Table S4. Statistical analysis of the three-chamber sociability test in $CRF_1$ receptor-deficient mice**. Statistical analysis of time (s) spent in the regions of interest (ROIs, side half-chambers) of the three-chamber apparatus and distance (m) travelled during the habituation and the sociability phases of the three-chamber test by male $CRF_1$+/+, $CRF_1$+/- and $CRF_1$-/- mice treated intraperitoneally (i.p.) with either saline or morphine (0.625 mg/kg). Genotype (G): $CRF_1$+/+ *vs*. $CRF_1$+/- *vs*. $CRF_1$-/-. Treatment (T): saline *vs*. morphine. Repeated measures (RM): mouse *vs*. object for the habituation and the sociability phase, habituation *vs*. sociability phase for distance. Further details are reported in the "statistical analysis" section of the manuscript.

| | **Habituation** | **Sociability** | **Distance** |
|---|---|---|---|
| **G** | $F_{2,58}$=2.528<br>P=0.089 | $F_{2,58}$=1.464<br>P=0.240 | $F_{2,58}$=2.280<br>P=0.111 |
| **T** | $F_{1,58}$=2.440<br>P=0.124 | $F_{1,58}$=0.002<br>P=0.963 | $F_{1,58}$=0.942<br>P=0.336 |
| **G x T** | $F_{2,58}$=5.069<br>P<0.01 | $F_{2,58}$=1.473<br>P=0.238 | $F_{2,58}$=1.525<br>P=0.226 |
| **RM** | $F_{1,58}$=0.023<br>P=0.879 | $F_{1,58}$=48.492<br>P<0.0001 | $F_{1,58}$=82.681<br>P<0.0001 |
| **G x RM** | $F_{2,58}$=0.019<br>P=0.981 | $F_{2,58}$=2.278<br>P=0.112 | $F_{2,58}$=1.772<br>P=0.179 |
| **T x RM** | $F_{1,58}$=0.031<br>P=0.861 | $F_{1,58}$=7.504<br>P<0.01 | $F_{1,58}$=2.000<br>P=0.163 |
| **G x T x RM** | $F_{2,58}$=0.083<br>P=0.920 | $F_{2,58}$=5.261<br>P<0.01 | $F_{2,58}$=5.145<br>P<0.01 |

**Table S5. Female $CRF_1$+/+ and $CRF_1$+/- mice fail to perform in the three-chamber task for sociability.** Number of female $CRF_1$+/+ and $CRF_1$+/- mice treated intraperitoneally (i.p.) with either saline or morphine (0.625 mg/kg) that visited both, only one or neither of the two side chambers of the three-chamber apparatus during the habituation phase (10 min) of the three-chamber test. Notably, only 2/8 saline-treated $CRF_1$+/+, 2/8 morphine-treated $CRF_1$+/+ and 3/8 morphine-treated $CRF_1$+/- female mice visited both side chambers of the apparatus. Thus, most of the animals tested met the exclusion criterion, i.e., exploration of each region of interest (ROI, side half-chamber) of the three-chamber apparatus for more than 80%, or less than 20%, of the total time spent in both ROIs during the habituation phase. This made impossible to assess morphine effects upon social behavior in female $CRF_1$ receptor-deficient mice using a reasonable number of animals.

| | **Side chambers visited** | | |
|---|---|---|---|
| **Genotype and treatment** | **Both** | **One** | **Neither** |
| **$CRF_1$+/+ saline (n=8)** | 2 | 2 | 4 |
| **$CRF_1$+/+ morphine (n=8)** | 2 | 3 | 3 |
| **$CRF_1$+/- saline (n=4)** | 4 | 0 | 0 |
| **$CRF_1$+/- morphine (n=8)** | 3 | 3 | 2 |

**Table S6. Statistical analysis of neuronal firing in C57BL/6J mice**. Statistical analysis of firing frequency (Hz) displayed by paraventricular nucleus of the hypothalamus (PVN) neurons expressing oxytocin (OXY) and/or arginine-vasopressin (AVP) in male and female C57BL/6J mice treated *per os* (p.o.) with either vehicle or antalarmin (20 mg/kg) and intraperitoneally (i.p.) with either saline or morphine (2.5 mg/kg). Pretreatment (P): vehicle *vs*. antalarmin. Treatment (T): saline *vs*. morphine. Further details are reported in the "statistical analysis" section of the manuscript.

| | **Male** | | **Female** | |
|---|---|---|---|---|
| | **OXY/AVP** | **AVP** | **OXY/AVP** | **OXY** |
| **P** | $F_{1,36}$=10.201 P<0.005 | $F_{1,45}$=0.340 P=0.562 | $F_{1,27}$=0.465 P=0.501 | $F_{1,34}$=0.094 p=0.761 |
| **T** | $F_{1,36}$=17.133 P<0.0005 | $F_{1,45}$=5.173 P<0.05 | $F_{1,27}$=13.685 P<0.001 | $F_{1,34}$=10.031 P<0.005 |
| **P x T** | $F_{1,36}$=10.186 P<0.005 | $F_{1,45}$=0.804 P=0.375 | $F_{1,27}$=0.006 P=0.939 | $F_{1,34}$=1.145 P=0.292 |
